# Interplay between stress and reproduction: Novel epigenetic markers in response to shearing patterns in Australian Merino sheep (*Ovis aries*)

**DOI:** 10.1101/2021.12.01.470842

**Authors:** Edward Narayan, Gregory Sawyer, Dylan Fox, Ryan Smith, Alan Tilbrook

## Abstract

In this study, we determined the effect(s) of shearing on Australian Merino ewes (*Ovis aries*). To test this research question, we used a suite of field and laboratory methods including GPS collars, wool cortisol and novel epigenetic markers identified using Illumina NovaSeq RRBS. Single shorn ewes (n =24) kept on their full fleece throughout the entire gestation period while twice shorn ewes (n =24) had their wool shorn early in gestation. We have discovered one locus (Chr20:50404014) which was significantly associated with different shearing treatments (twice or single shorn ewes), (FDR = 0.005). This locus is upstream of a protein coding gene (ENSOARG00000002778.1), which shows similarities to the forkhead box C1 (FOXC1) mRNA using BLAST searches. We discovered that 36 gene loci were significantly modulated either between different shearing treatments or late vs early pregnancy ewes. Similarly, in lambs we identified 16 annotated gene loci that were significant between late vs early pregnancy. Early shorn ewes grazed 10% higher and maintained stronger body condition. Wool cortisol levels were significantly lower in the early shorn ewes during mid- and late gestation. Lambs bred from twice shorn ewes had on average better visual wool quality parameters in terms of micron, spin finesses and curvature. Collectively, this research provides a new dataset combining physiological, molecular epigenetics and digital tracking indices that advances our understanding of how Merino ewes respond to shearing frequency and this information could guide further research on sheep breeding and welfare.

**Significance Statement:** This study provides novel data on the molecular epigenetic signatures in Merino sheep under exposure to natural environmental and management factors. We have discovered DNA methylation profiles from ewes and lambs that are directly associated with whole- animal physiology, development and growth. The baseline data can provide a useful resource for future research in many key areas such as animal welfare, diseases and climatic resilience that will benefit sheep and wool production.

## Introduction

In the past decade, there has been an increase in scientific reporting into the effects of adversity in early life on the participant’s DNA profile within both human and animal studies (1, 2). This emerging research area continues to be driven by scientists globally to better understand a wide range of effects caused by a variety of intrinsic and extrinsic influences on the DNA profile of ongoing generations within observed genotypes. From within the cell of the early stage developing embryo, transcriptional and epigenetic changes to the cell are occurring *via* remodelling and reprogramming within the cell nucleus. What is still unclear is if/how external factors such as management practices could alter DNA during critical life-history phases such as reproduction. This science is what is known as epigenetics (3). Generally, epigenetics represents the genome-wide study of the distribution of methylated and unmethylated nucleoside residues within the genome (4). (5) concluded that epigenetics refers to effects on phenotype (or on patterns of gene expression) that are passed from one generation to the next by molecules in the germ cells and that cannot be explained by Mendelian genetics (or by changes to the primary DNA sequence). In production animal research, recently it has been highlighted the need of epigenetic evaluation of maternal-fetal nexus especially in relation to environmental factors (e.g. climate), nutrition and post-natal development and growth of progeny (6).

Early biomedical studies into animals’ epigenetics have been focused on mice due to their ability to reproduce quickly and for researchers to gain fast results with multiple offspring from the same female. Due to the nature of sheep growth and time to reach puberty, a predominant single offspring and the lack of funding for epigenetic research into the sheep there have been very limited research into this field (see 7,8). However, due to foresight by early researchers there have been substantial advances in current genomic technologies to allow for development of genome analysis and sequencing in livestock (9). To our knowledge, there is no peer reviewed scientific literature of the epigenetic effect on the sheep’ DNA caused by a basic management intervention such as shearing. Our earlier research showed that embryos from the same sire and dam that were placed into surrogates and raised under the same environmental and management regimes had significant varying phenotypic (wool quality trait) outcomes (10, 11). Maternal stress at the early stages of pregnancy appears to result in greater implication of the epigenetic changes than that trigged post-parturition (12). The placenta acts as a connection between the mother and the developing fetus and stress activates the maternal hypothalamo-pituitary adrenal (HPA) axis and triggers stress hormone or glucocorticoid (GC) synthesis that reaches the foetus by transplacental passage (13). Prenatal stress and prenatal exposure to GCs have been shown to have long-term effects on the expression of numerous genes associated with HPA function, neurologic function and phenotype (14). In sheep, latest research has demonstrated that pharmacologically elevated cortisol can also result in negative effects on the animal’s body condition and phenotype (wool fibre diameter) [15].

Shearing is a well validated management intervention and known to generate acute stress responses in sheep (16). Our interest was to monitor the maternal ewe’s HPA-axis activity between shearing treatments; therefore, we employed a well-established wool cortisol assay (17, 18). Benefits of early or even mid-pregnancy shearing is well known across various breeds of sheep such as increase in shelter seeking, cleaner udder areas with lesser wool, resulting in lambs born heavier and with improved survival chances (19). However, there has been no previous studies conducted in sheep on the influence of shearing pattern (once a year verse twice a year) on maternal stress physiology, grazing behaviour and epigenetic effects on lamb phenotype. We hypothesised that shearing frequency (twice shorn versus single shorn) could influence the grazing activity of pregnant ewes with underlying changes to molecular epigenetic profiles of ewes, wool cortisol and body condition, with flow-on positive benefits of early pregnancy shearing on lamb phenotype (wool quality).

## Results

### Epigenetic DNA Methylation

Genewise negative binomial generalized linear models (glmLRT) were used to identify differences in methylation between different shearing treatments and pregnancy durations. In total 22 samples (out of 24 ewes) with one shearing treatment and 23 (out of 24 ewes) with two shearing treatments were collected in this study (1 unknown). Moreover, 7 and 27 samples were collected from sheep with a short and long pregnancy (12 unknown), respectively.

One locus (Chr20:50404014) was significantly associated with different shearing treatments (twice or single shorn ewes), (FDR = 0.005). This locus is upstream of a protein coding gene (ENSOARG00000002778.1), which shows similarities to the forkhead box C1 (FOXC1) mRNA using BLAST searches. No significant differences were measured for pregnancy duration (early or late) using glmLRT (FDR > 0.134). A MDS plot did not separate the ewe samples according to pregnancy duration or shearing treatment (Fig. 1).

**Figure 1.**
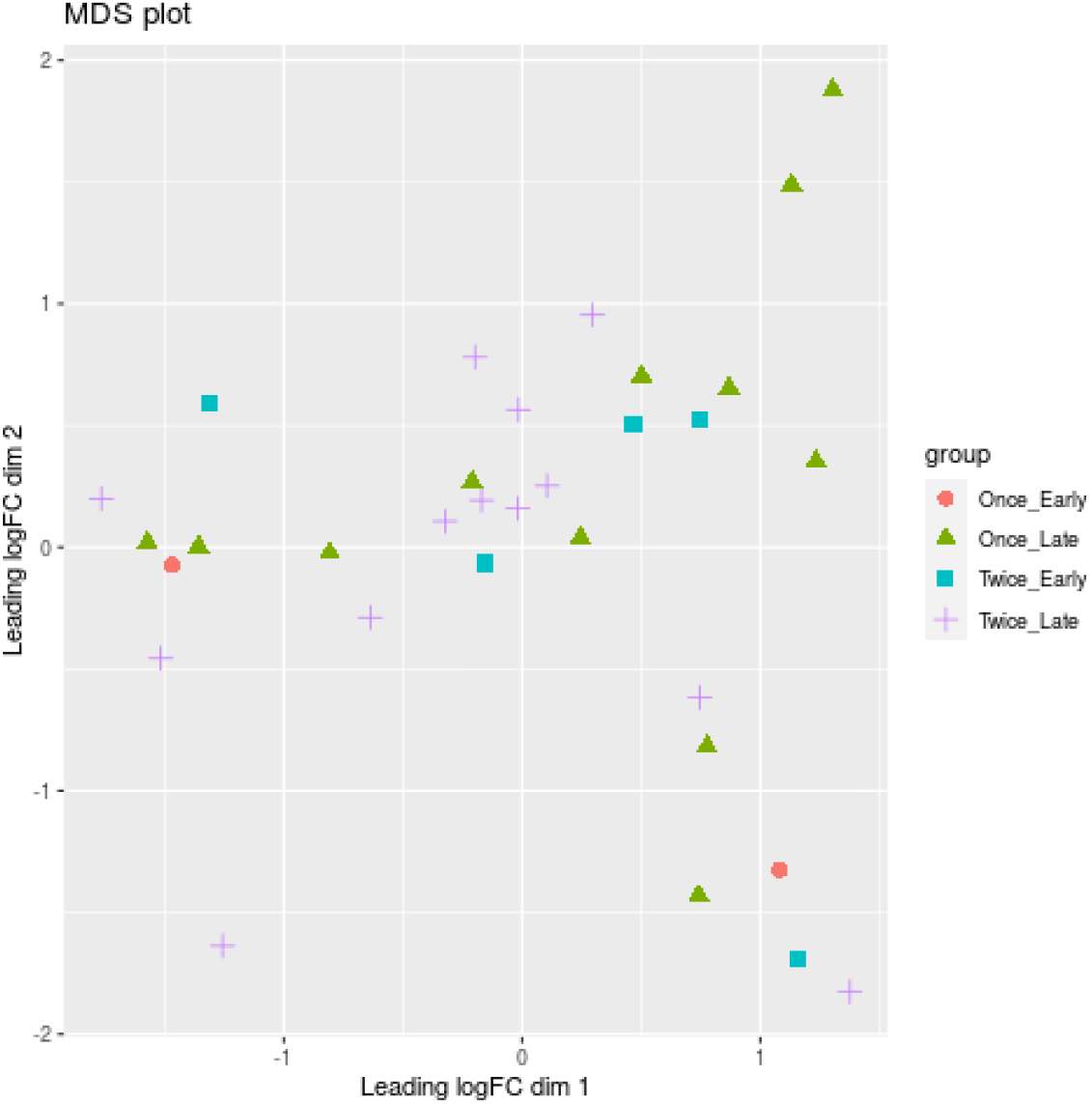
Multidimensional scaling (MDS) plot based on all methylation sites. Each dot represents an ewe sample and is colored by the pregnancy duration (early vs late) and shearing treatment (once vs twice). Only the 50 most significant loci were used for the MDS analysis.

Differential methylation analysis was repeated using a combination of the shearing treatment and pregnancy status. The comparisons (1) Late pregnancy vs early pregnancy for sheep with one shearing treatment and (2) Late pregnancy vs early pregnancy for sheep with two shearing treatments were carried out to identify associations between loci and pregnancy duration for sheep with either one or two shearing events.

As shown in figure 2.0A, the Venn diagram illustrates the loci that were up/down in both or in each comparison (1) and (2). 8 loci are significantly upregulated while 6 loci were downregulated between late vs early pregnancy in sheep that were sheared once or twice. The annotation of these loci is provided as a bar graph in Fig. 3.0A.

**Figure 2.0.**
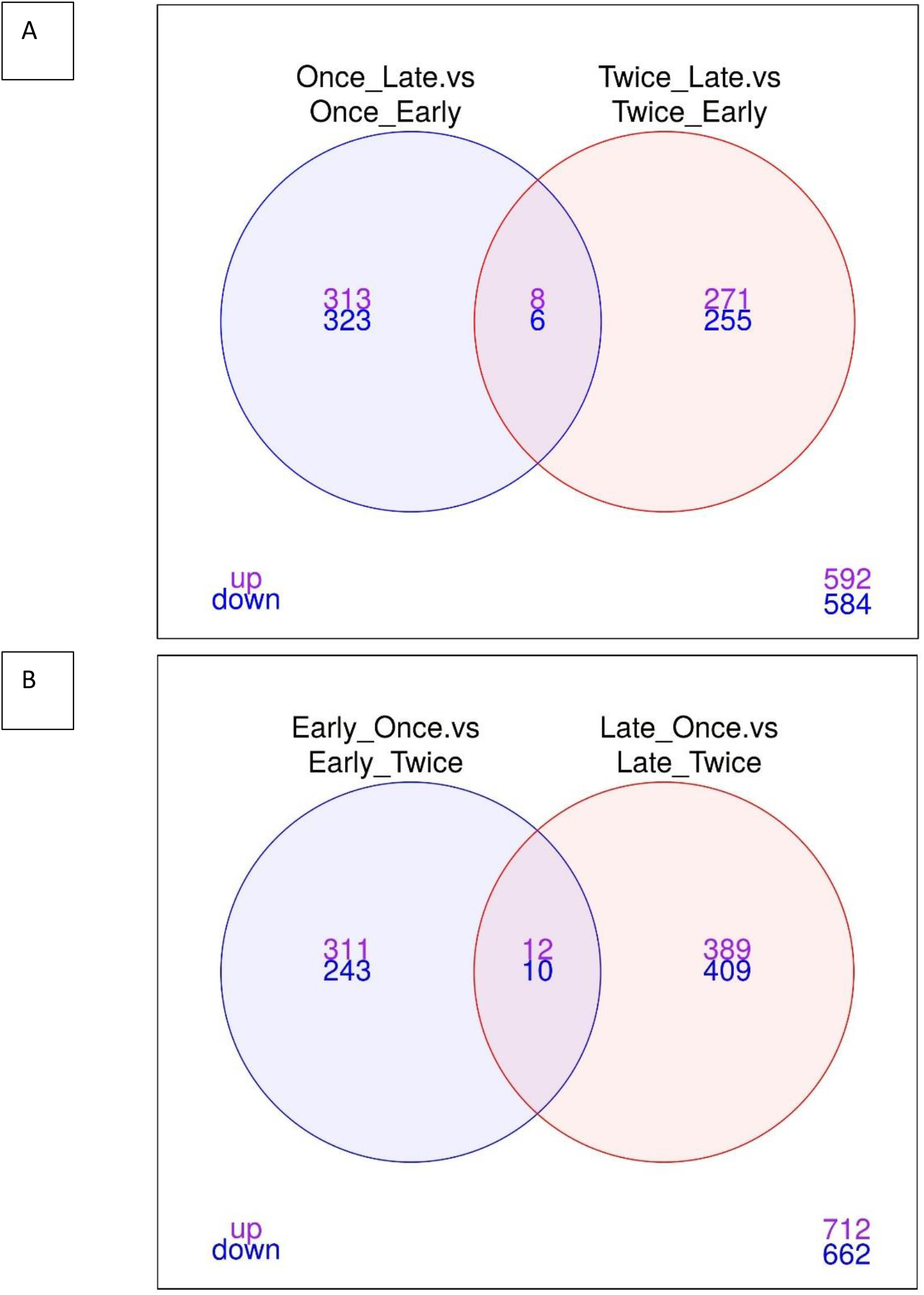
Venn Diagram illustrating the up and down-regulation of loci between different comparisons. The venn diagrams illustrate the core genes for (A) “Once sheared and late pregnancy vs once sheared and early pregnancy” as well as in “Twice sheared and late pregnancy vs twice sheared and early pregnancy”. (B) Early pregnancy and once sheared vs early pregnancy and twice sheared as well as late pregnancy and once sheared vs late pregnancy and twice sheared sheep. Only features with a FDR < 0.05 and logFC > 0 were included.

The comparisons (3) Once vs twice sheared for sheep with early pregnancy and (4) Once vs twice sheared for sheep with late pregnancy were calculated to identify associations between loci and shearing treatment for sheep with either early or late pregnancy. The core loci, ie loci that are either significantly up- or downregulated between different shearing treatments for sheep with either pregnancy duration are visualized in a venn diagram (Figure 2B) and bargraphs (Figure 3B). In total 12 loci were upregulated and 10 downregulated.

**Figure 3.0.**
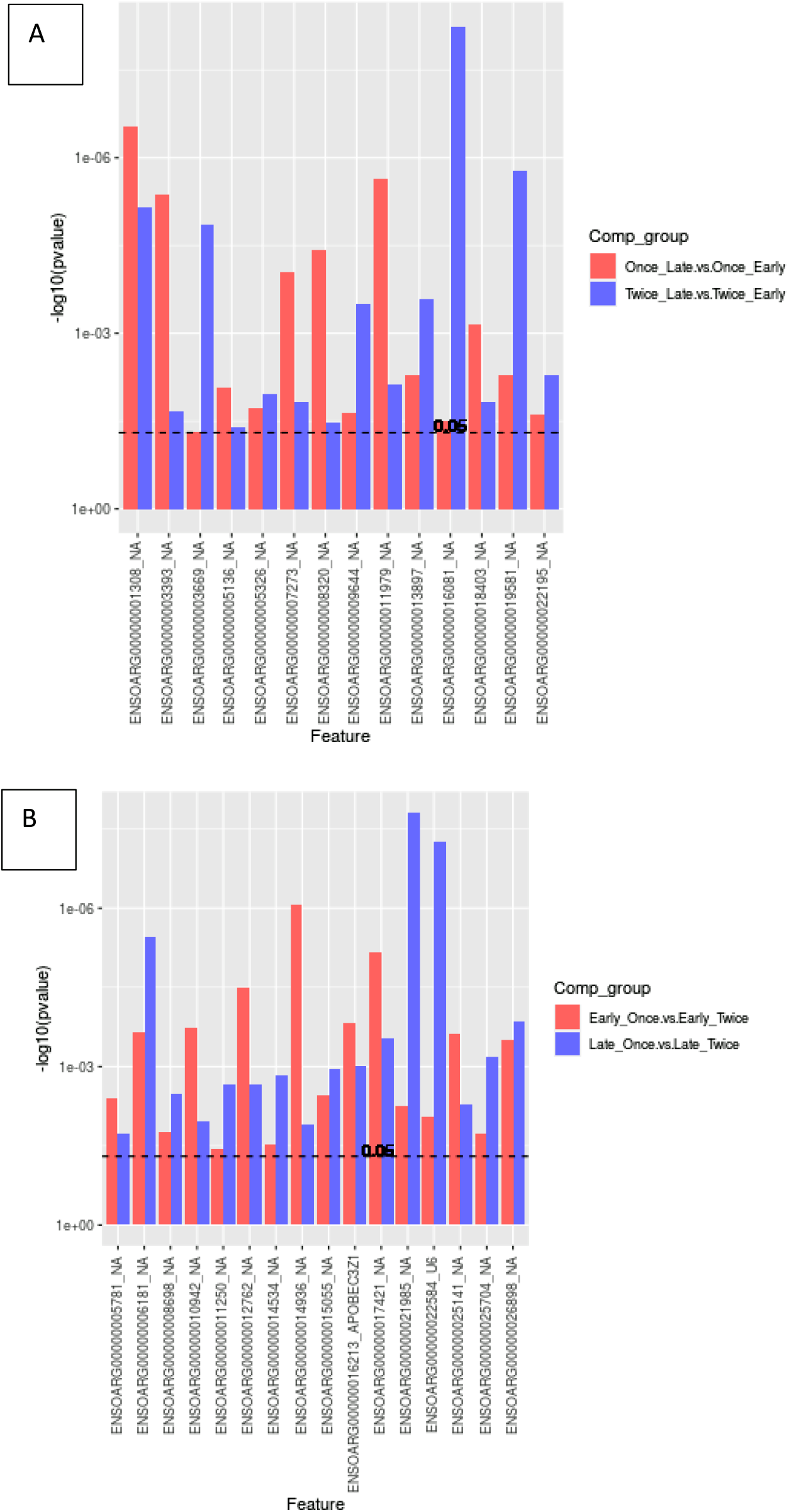
bargraph plot showing the annotations and significantly different overlapping methylation sites (in reference to the Venn diagrams, Fig2 A-B). (A) A total of 14 (8/6) loci were up- or down-regulated for comparisons of pregnancy scanning for once or twice sheared sheep. All loci have not a gene symbol (NA, not available). (B). In total 22 (12/10) loci were up- or down-regulated between shearing treatment including APOBECC3Z1 gene and U6.

Volcano Plot

**Figure 4.0.**
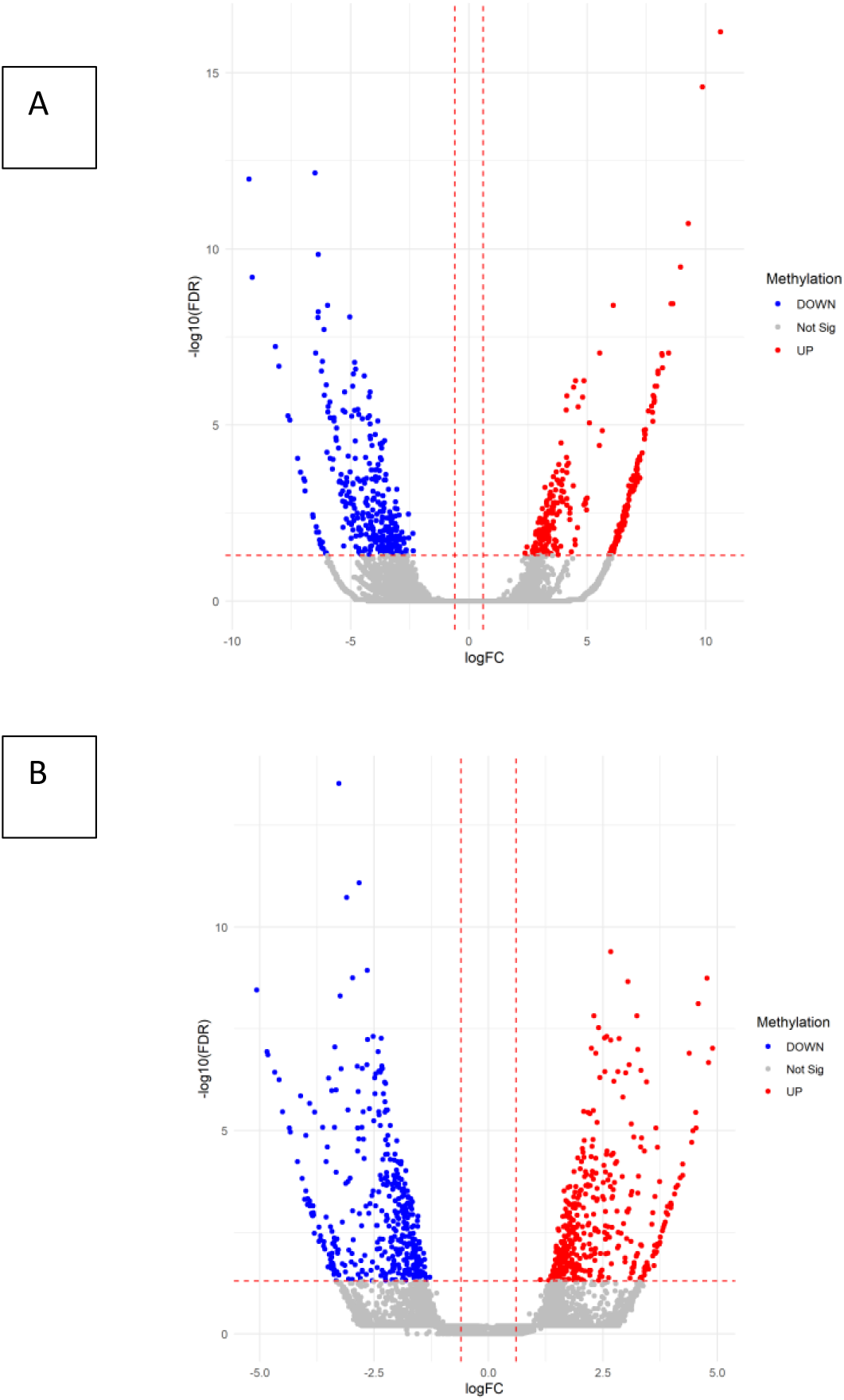
Volcano plots summarises how many loci were significant and how different the log fold change was for shearing treatment (A) or pregnancy scanning (B).

**Table 1.0.**
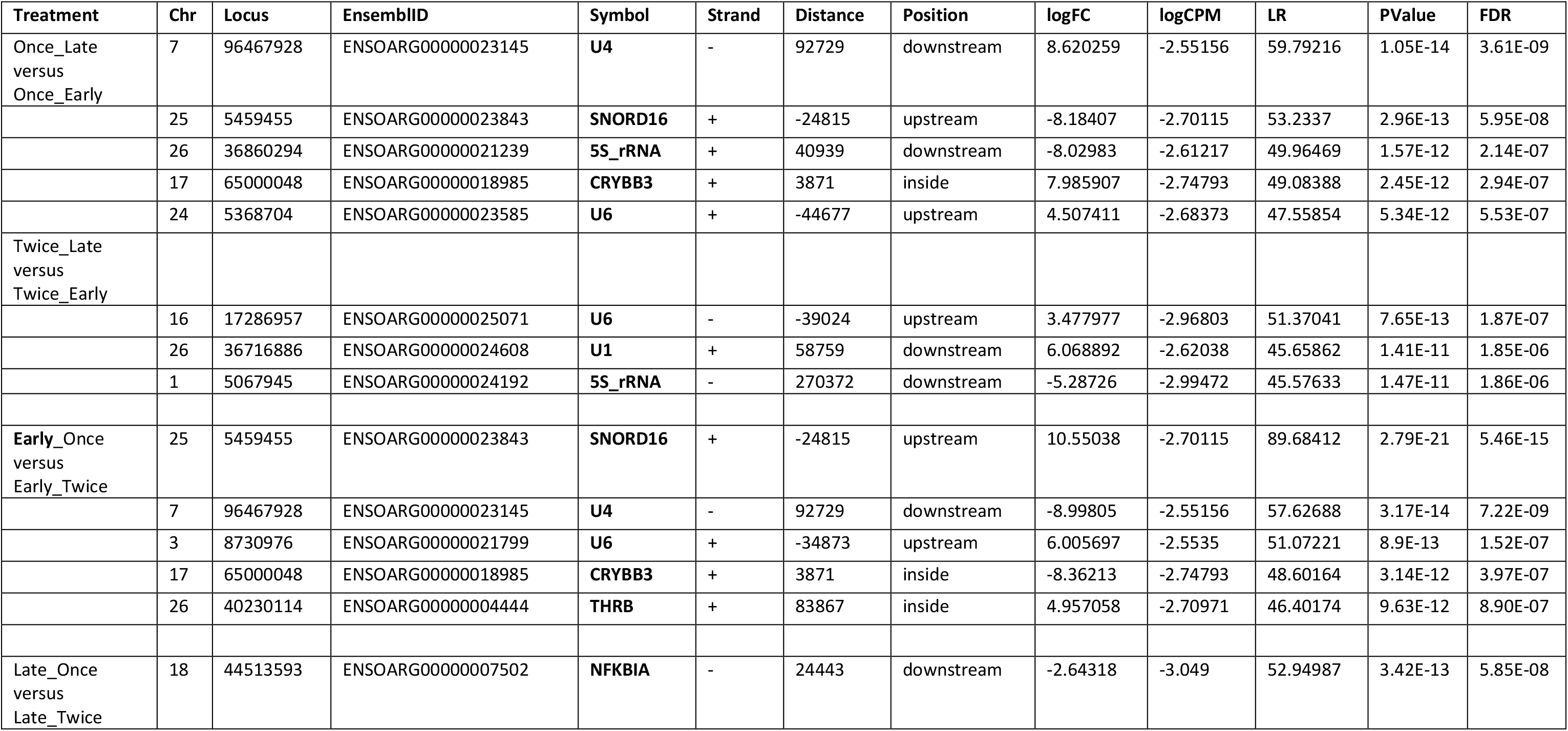
Statistically significant gene loci (with known annotation) in Merino ewes for shearing treatment and pregnancy scanning.

### Ewe Body Condition Scores

Ewes from both treatments (shearing groups) showed variability in their body condition score. However, the twice shorn ewes maintained their body condition during mid- and late pregnancy on average higher than the once only shorn ewes (Table 2, p < 0.05). Average range of condition scores (2.5-3.5) is typical of healthy ewes.

**Table 2.** Count, mean, and standard error values for ewe condition score across once and twice shorn treatments. Means and standard errors are rounded to two decimal places where necessary.

| **Pregnancy**<br>**status** | **Once Shorn** |  |  |  | **Twice Shorn** |  |  |  |
| --- | --- | --- | --- | --- | --- | --- | --- | --- |
|  | *n* | Median | Mean | Standard<br>Error | *n* | Median | Mean | Standard<br>Error |
| **Mid** | 24 | 2.6 | 2.61 | 0.0781 | 24 | 2.75 | 2.77 | 0.0721 |
| **Late** | 24 | 2.5 | 2.54 | 0.1009 | 24 | 3 | 2.77 | 0.1369 |
| **Post-**<br>**lambing** | 24 | 1.5 | 1.5 | 0.0630 | 24 | 1.5 | 1.48 | 0.0923 |

### Wool Cortisol Profiles

Mean Wool Cortisol Metabolites (WCM) of Merino Ewes initially decreased between pre-joining and mid-pregnancy, however increased between April (mid gestation) and May (late gestation) across both treatments. However, once shorn ewes recorded the highest average WCM results (Fig. 2, p < 0.05).

### Grazing Activity

**Figure 5.**
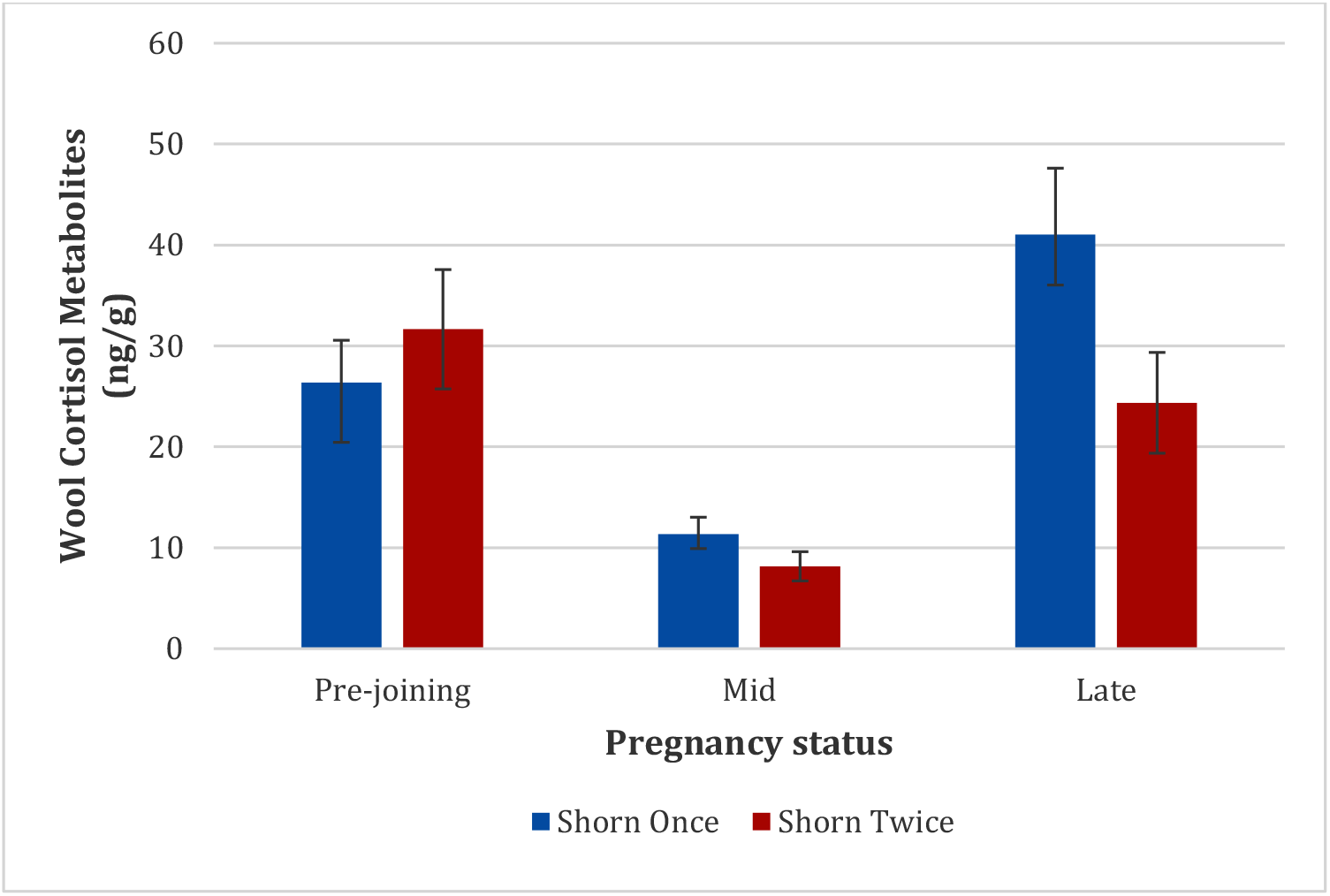
Mean Wool Cortisol Metabolites (WCM) of Merino Ewes Shorn Once and Twice. Error bars represent standard error of individual data set. N = 24 per group.

Collar data were analyzed between post-shearing (14 April, 2019) and 24 May, 2019 (late-gestation) to determine where shearing once (n = 9 sheep with collar data) or twice (n = 11 sheep with collar data) led to a significance change in grazing activity between the treatments (p <0.05, Fig. 3). Statistical analysis showed a significant difference in grazing frequency between once and twice shorn ewes (p < 0.05). Twice shorn sheep were grazing proportionally more than once shorn sheep by >10% during mid-late gestation.

**Figure 6.**
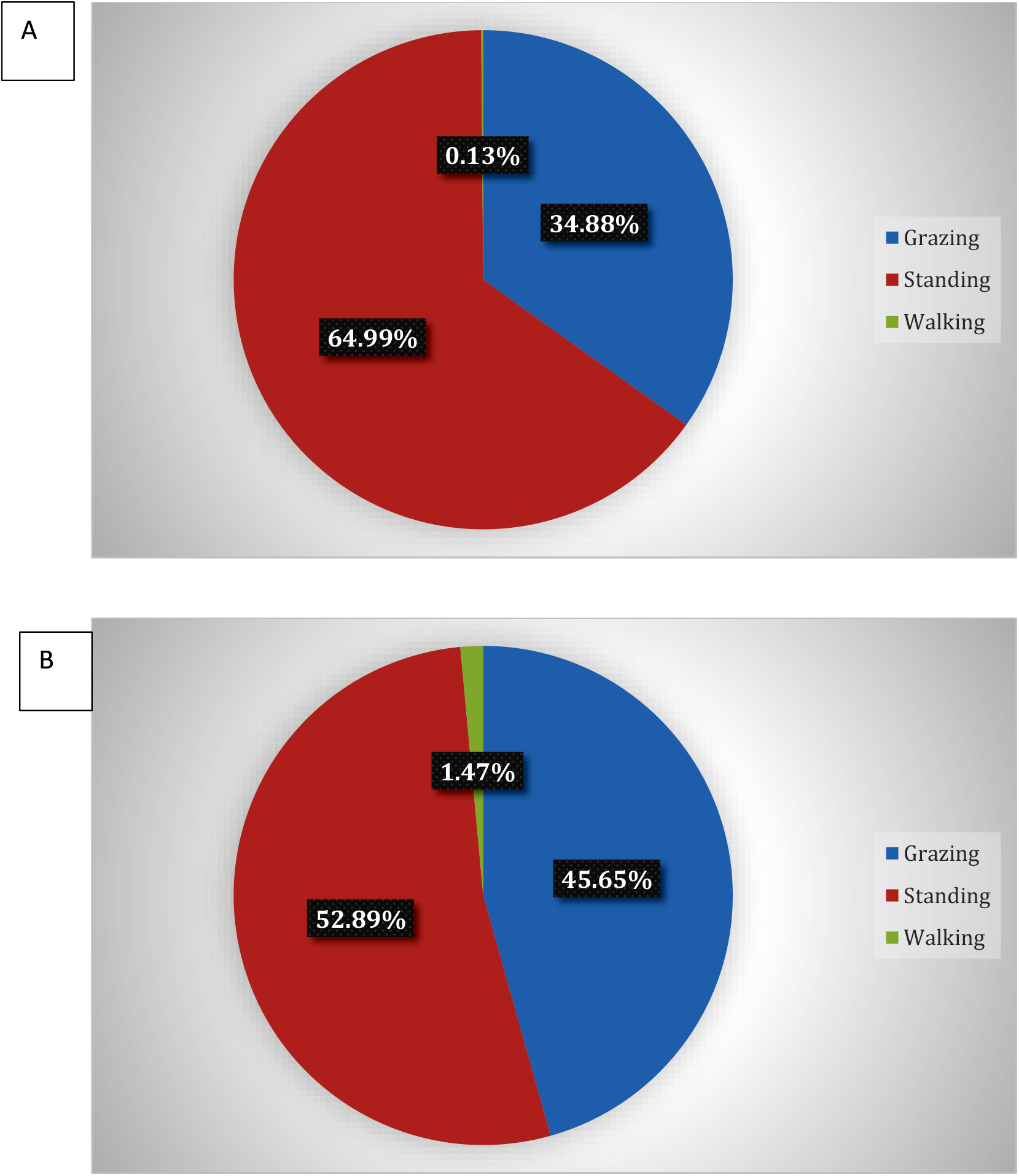
A: Mean (*n*= 9) activity budget of Merino ewes shorn once. B: Mean (*n*= 11) activity budget of Merino ewes shorn twice.

### Lamb phenotype and molecular data

The results in the table below show wool characteristics of the lambs that were matched to their ewes using the DNA parentage test. The lambs from the shorn twice cohort of ewes had a visually finer wool (Average micron column), higher average comfort factor of 0.4 percent (Ave COMFF column) and spinning fineness difference between shearing frequency groups was 0.9 microns (Ave Spin Fineness Column).

**Table 3.**
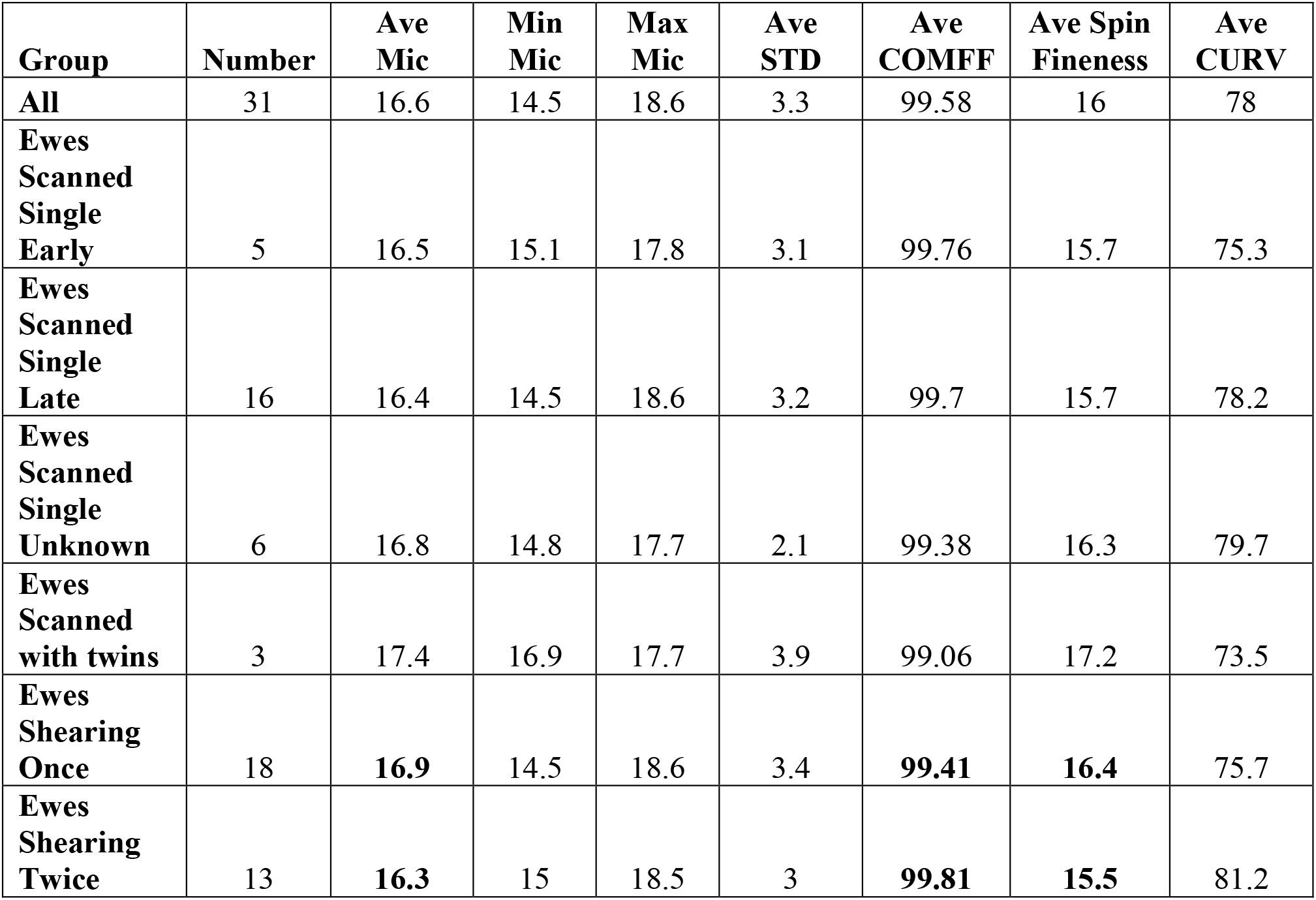
Average raw wool micron, CC%, comfort factor average spinning fineness or average curvature of lambs grouped to ewes (Group column).

| **Group** | **Number** | **Ave Mic** | **Min Mic** | **Max Mic** | **Ave STD** | **Ave COMFF** | **Ave Spin Fineness** | **Ave CURV** |
| --- | --- | --- | --- | --- | --- | --- | --- | --- |
| **All** | 31 | 16.6 | 14.5 | 18.6 | 3.3 | 99.58 | 16 | 78 |
| **Ewes Scanned Single Early** | 5 | 16.5 | 15.1 | 17.8 | 3.1 | 99.76 | 15.7 | 75.3 |
| **Ewes Scanned Single Late** | 16 | 16.4 | 14.5 | 18.6 | 3.2 | 99.7 | 15.7 | 78.2 |
| **Ewes Scanned Single Unknown** | 6 | 16.8 | 14.8 | 17.7 | 2.1 | 99.38 | 16.3 | 79.7 |
| **Ewes Scanned with twins** | 3 | 17.4 | 16.9 | 17.7 | 3.9 | 99.06 | 17.2 | 73.5 |
| **Ewes Shearing Once** | 18 | **16.9** | 14.5 | 18.6 | 3.4 | **99.41** | **16.4** | 75.7 |
| **Ewes Shearing Twice** | 13 | **16.3** | 15 | 18.5 | 3 | **99.81** | **15.5** | 81.2 |

A MDS plot did not separate the ewe samples according to pregnancy duration or shearing treatment (Fig. 7A-B).

**Figure 7.**
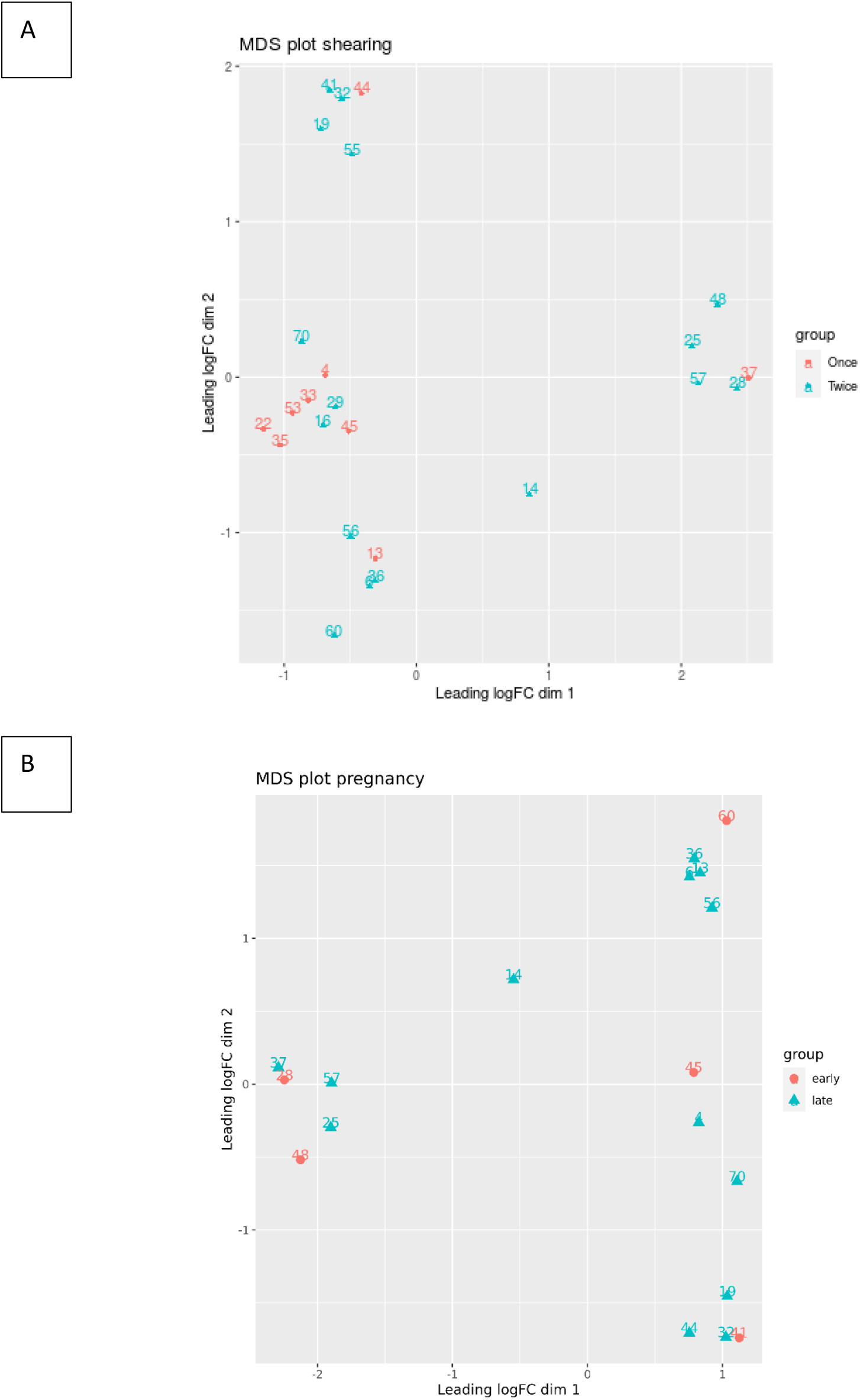
Multidimensional scaling (MDS) plot based on all methylation sites. Each dot represents a lamb sample and is colored by the pregnancy duration (early vs late) and shearing treatment (once vs twice). Only the 50 most significant loci were used for the MDS analysis.

**Table 4.0.** Statistically significant gene loci (with known annotation) in Merino lambs for shearing treatment and pregnancy scanning.

| Chr | Locus | feature | feature_strand | distancetoFeature | insideFeature | external_gene_name | logFC | logCPM | LR | PValue | FDR |
| --- | --- | --- | --- | --- | --- | --- | --- | --- | --- | --- | --- |
| 18 | 31863214 | ENSOARG000000021220 | - | -22750 | upstream | 5S_rRNA | 6.063927 | -3.57787 | 42.24772 | 8.04E-11 | 1.43E-05 |
| 3 | 910106 | ENSOARG000000005761 | - | -103221 | upstream | IGFBP7 | -3.87275 | -3.5854 | 37.6132 | 8.63E-10 | 9.76E-05 |
| 8 | 74436878 | ENSOARG000000022287 | + | -29373 | upstream | 5S_rRNA | -5.02242 | -3.06516 | 36.21928 | 1.76E-09 | 0.000164 |
| 1 | 11654566 | ENSOARG000000021462 | + | -30339 | upstream | oar-mir-29b-1 | -3.04434 | -3.46646 | 34.62464 | 4.00E-09 | 0.000299 |
| 20 | 47000448 | ENSOARG000000024332 | + | 9534 | downstream | U6 | 5.634211 | -3.27042 | 34.56958 | 4.11E-09 | 0.000303 |
| 10 | 19227897 | ENSOARG000000006000 | + | -170 | upstream | B4GALT7 | 5.58518 | -3.2474 | 33.0828 | 8.83E-09 | 0.000561 |
| 1 | 2.41E+08 | ENSOARG000000021546 | + | 7002 | downstream | SNORA70 | 3.196796 | -2.39691 | 32.28392 | 1.33E-08 | 0.000782 |
| 26 | 471140 | ENSOARG000000008561 | - | -3715 | upstream | UCP2 | -2.60014 | -3.26207 | 30.99755 | 2.58E-08 | 0.001317 |
| 1 | 9519659 | ENSOARG000000013636 | + | -13666 | upstream | CDKN1A | 5.424016 | -3.66533 | 30.32599 | 3.65E-08 | 0.001645 |
| 4 | 1.18E+08 | ENSOARG000000024658 | + | -32194 | upstream | SNORD14 | 2.788125 | -3.14698 | 30.27497 | 3.75E-08 | 0.001669 |
| 21 | 47754634 | ENSOARG000000010362 | + | -5544 | upstream | MRCL3 | 4.345068 | -3.27269 | 29.57888 | 5.37E-08 | 0.002147 |
| 26 | 37758950 | ENSOARG000000011711 | + | -275 | upstream | GATD3A | -2.55053 | -3.51928 | 27.61278 | 1.48E-07 | 0.00483 |
| 3 | 2208581 | ENSOARG000000013752 | + | -12598 | upstream | NOG | -2.81314 | -2.53443 | 27.4112 | 1.64E-07 | 0.005191 |
| 20 | 27480246 | ENSOARG000000005005 | + | -1276 | upstream | NPAS4 | 2.641856 | -3.12799 | 26.54263 | 2.58E-07 | 0.007035 |
| 1 | 86523392 | ENSOARG000000023803 | - | 15090 | downstream | 7SK | 2.744371 | -2.87492 | 26.2391 | 3.02E-07 | 0.007862 |
| 22 | 19864886 | ENSOARG000000012855 | + | 14588 | downstream | ST3GAL4 | -4.22223 | -2.92788 | 24.77667 | 6.44E-07 | 0.013636 |
| 17 | 72105115 | ENSOARG000000011166 | + | 17243 | downstream | BLOC1S4 | -2.31982 | -3.31215 | 23.95507 | 9.86E-07 | 0.019141 |
| 8 | 89756571 | ENSOARG000000018593 | - | -386 | upstream | HADHA | -4.17315 | -3.3325 | 23.31672 | 1.37E-06 | 0.024997 |
| 13 | 39650849 | ENSOARG000000008334 | + | 100280 | downstream | VEGFA | -2.46058 | -3.41285 | 22.33421 | 2.29E-06 | 0.037422 |
| 3 | 1.36E+08 | ENSOARG000000019100 | + | 7808 | downstream | ADIG | 2.320133 | -3.5414 | 22.08821 | 2.60E-06 | 0.040812 |
| 26 | 636096 | ENSOARG00000012211 | - | 68 | inside | RPS25 | 2.385044 | -3.43847 | 21.86493 | 2.93E-06 | 0.044788 |
| 9 | 80656245 | ENSOARG00000020177 | + | 585 | inside | PDIA5 | 4.708262 | -2.50831 | 21.54701 | 3.45E-06 | 0.050687 |
| 16 | 4599653 | ENSOARG00000010800 | + | -231 | upstream | PRDX2 | -2.34846 | -3.22251 | 21.36714 | 3.79E-06 | 0.054634 |

## Discussion

This research aimed to study the molecular epigenetics, stress physiology and behaviour of pregnant Merino ewes using a combination of field and lab based methods to test the original hypothesis that shearing frequency (twice shorn versus single shorn) could influence the grazing activity of pregnant ewes with underlying changes to molecular epigenetic profiles of the ewes, wool cortisol and body condition, with flow-on positive benefits of early pregnancy shearing on lamb phenotype (wool quality).

Based on the results from the experiment, it was determined that shearing frequency did result in significant molecular epigenetic change in the pregnant ewes. We discovered one locus (Chr20:50404014) which was significantly associated with different shearing treatments (twice or single shorn ewes), (FDR = 0.005). This locus is upstream of a protein coding gene (ENSOARG00000002778.1), which shows similarities to the forkhead box C1 (FOXC1) mRNA using BLAST searches. There was lower methylation level in Once versus Twice shorn ewes, which indicated the locus is downregulated. FOXC1 gene provides instructions for making a protein that attaches (binds) to specific regions of DNA and regulates the activity of other genes. On the basis of this action, the FOXC1 protein is called a transcription factor. The FOXC1 protein plays a critical role in early development of the eye, normal development of heart, kidneys and brain. It also plays critical role in adult management of oxidative stress associated with the visual cortex. The FOXC1 gene has been identified in Chinese native sheep breeds in a recent genomic profiling study, as a novel series of vision-related genes [20]. The FOXC1 gene belongs to forkhead family of transcription factors characterized by a distinct DNA-binding forkhead domain. These transcription factors are involved in regulating embryonic and ocular development and FOXC1 mutations are associated with various glaucoma phenotypes, including primary congenital glaucoma and Axenfeld–Rieger syndrome, which characterized by specific ocular anomalies [21]. To our knowledge, our research is the first to demonstrate the epigenetic modulation of FOXC1 gene during pregnancy in Merino sheep in a natural grazing study, and this should be explored further in relation to the consequences on embryonic and ocular development of Merino lambs.

We found no significant differences were measured for pregnancy duration (early or late) using glmLRT (FDR > 0.134). One explanation may be loss of power. By separating the data into early and late, we reduced the number of samples for the comparisons. A MDS plot did not separate the ewe samples according to pregnancy duration or shearing treatment. Differential methylation analysis was repeated using a combination of the shearing treatment and pregnancy status to identify associations between loci and pregnancy or shearing events. We discovered that 8 loci were significantly upregulated while 6 loci were downregulated between late vs early pregnancy in sheep that were sheared once or twice (all loci are novel and not previously annotated with BLAST search). Secondly, in total 12 loci were upregulated and 10 downregulated between different shearing treatments for sheep with either pregnancy duration.

The annotated epigenetically modulated genes for Merino ewes as determined using gene cards (see table 1) included the following genes. APOBEC3 gene (this is Apolipoprotein B MRNA Editing Enzyme Catalytic Subunit). U1, U4, U6 - RNA gene affiliated with the snRNA class. Among its related pathways are mRNA Splicing - major pathway and RNA transport. U6, which is a type of small nuclear RNA (snRNA) and is highly conserved among species. U6 snRNA located at the heart of the spliceosome participates in the processing of mRNA precursors. U6 is very stable because of the combination of small nuclear ribonucleoprotein complexes, a 5′ cap, a 3′U-rich tail, and the capacity for self-and/or U4 hybridization. The half-life value is approximately 24 hours. U6 is one of the most widely used internal reference genes for miRNA. U6 has been used as an internal reference gene in renal tissue, cell lines and peripheral blood mononuclear cells in kidney disease patients (see: 22). SNORD16 - (Small Nucleolar RNA, C/D Box 16) is an RNA Gene, and is affiliated with the snoRNA class. 5S_rRNA - an integral component of the large ribosomal subunit in all known organisms with the exception only of mitochondrial ribosomes of fungi and animals. It is thought to enhance protein synthesis by stabilization of a ribosome structure. CRYBB3 - is a Protein Coding gene. Diseases associated with CRYBB3 include Cataract 22, Multiple Types and Cataract 24. Gene Ontology (GO) annotations related to this gene include structural constituent of eye lens.

An important paralog of this gene is CRYBB2. THRB - The protein encoded by this gene is a nuclear hormone receptor for triiodothyronine. It is one of the several receptors for thyroid hormone, and has been shown to mediate the biological activities of thyroid hormone. Knockout studies in mice suggest that the different receptors, while having certain extent of redundancy, may mediate different functions of thyroid hormone. Mutations in this gene are known to be a cause of generalized thyroid hormone resistance (GTHR), a syndrome characterized by goiter and high levels of circulating thyroid hormone (T3-T4), with normal or slightly elevated thyroid stimulating hormone (TSH). NFKBIA - This gene encodes a member of the NF-kappa-B inhibitor family, which contain multiple ankrin repeat domains. The encoded protein interacts with REL dimers to inhibit NF-kappa-B/REL complexes which are involved in inflammatory responses.

The annotated epigenetically modulated genes for Merino lambs as determined using gene cards (see table 1) included the following genes; 5S_rRNA; IGFBP7; oar-mir-29b-1, U6, B4GALT7, SNORA70, UCP2, CDKN1A, SNORD14, MRCL3, GATD3A, NOG, NPAS4, 7SK, ST3GAL4, BLOC1S4, HADHA, VEGFA, ADIG, RPS25, PDIA5 and PRDX2 (see table 2.0). Some of these interesting candidate genes are discussed.

Insulin-like growth factor-binding protein 7 (IGFBP7) is an interesting gene relevant to pigmentation in merino sheep and located within a 3 Mb window of ovine chromosome 6 (OAR6), in a region that also contains the KIT gene to which piebald was first associated [23]. IGFBP7 could be studied further to boost molecular markers for the elimination of contamination of white wool with contaminated fibres. This is possible with availability of IGFBP7 transcripts now available for sheep [24]. NOG gene has earlier been identified as one of the candidate inhibitor genes of secondary follicle differentiation in developing Merino fetus in utero [25]. 5S rRNA gene is involved in ribosome functioning [26]. SNORA70 gene is important in relation to feed intake, body composition and live weight and earlier study based on the local adaptation of Creole cattle genome diversity in the tropics, reported an association between SNORA70 populations adapted either to cold or heat conditions, indicating its importance for metabolic adaptation in both thermal conditions [27; 28]. Our results for epigenetic modulation of SNORA70 gene also confirms the differential expression work done earlier by [28], whereby maternal under- and overnutrition during gestation in Merino ewes affected small RNA species in the offspring lamb including SNORD113 and SNORA70. SNORD14 is highly significant in relation to heat stress [29]. These are small, nuclear, noncoding RNA that are responsible for guiding RNA for post-transcriptional modifications. Further research is warranted to understand how access to adequate maternal nutrition during gestation impacts of fetal development. MicroRNA such as oar-mir-29b-1 is a non coding RNA that has been shown to play a major role in neurological brain development in rodents and human models through its influence on the expression of DNA methyltransferase *Dnmt3a,* which is responsible for the catalyzation of CH methylation. Methyl-CH (mCH) marks have been hypothesized to regulate neuronal diversity by fine-tuning gene transcription across the genome (30). B4GALT7, beta-1,4- galactosyltransferase 7 has a key role in collagen-network formation. B4GALT7 is involved in glycosylation, has a role in connective-tissue disorders and is related to disturbed fibril organisation and proteoglycan synthesis. B4GALT7 is highly expressed in the growth plate, especially in the proliferative zone, mutations have been shown to cause dwarfism in livestock e.g. horses (31). Uncoupling protein (UCP) 2 is a widely expressed mitochondrial protein whose precise function is still unclear but has been linked to mitochondria-derived reactive oxygen species production. Thus, the chronic absence of UCP2 has the potential to promote persistent reactive oxygen species accumulation and an oxidative stress response as well as impaired glucose stimulated insulin production (32). Myosin regulatory subunit (MRLC3) plays an important role in regulation of both smooth muscle and nonmuscle cell contractile activity *via* its phosphorylation. Implicated in cytokinesis, receptor capping, and cell locomotion. MRLC2 expression has been studied in skeletal muscle contraction of goats (33). Neuronal PAS domain-containing protein 4 (Npas4), a member of the PAS family characterized by a conserved basic-helix-loop-helix motif and PAS domain, acts as an inducible immediate early gene (IEG) activated with minutes of stimulation to regulate the formation of inhibitory synapses. Importantly, the physicochemical properties and reveals the neuro-modulatory role of Npas4 in crucial pathways involved in neuronal survival and neural signaling (34). In a context relevant to future research in livestock welfare, studies in rodent model have shown changes in Npas4 expression levels in response to stress such as fear, sleep deprivation, anxiogenic environments, maternal separation and absence of Npas4 has been associated with higher expression levels of cytokines IL-6 and TNF-α and modulation of brain- circuits important for cognitive and emotional sense (34). ADIG is also known to be a lipid-producing protein is a novel transcription factor for regulating and controlling the differentiation and proliferation of adipocytes. The sequence of the ADIG gene and protein of human, mouse, rat, pig and cattle is highly conserved, and it has similar regulatory and biological functions in the mammalian body. ADIG is a newly focused transcription factor in the field of animal husbandry, which regulates the expression of PPARc and participates in the regulation and control of the production of fat cells. Its closely associated with glycolipid metabolism and sugar metabolism and targeted regulation of ADIG could be applied to better understand the mechanism of IMF deposition in livestock (35). Peroxiredoxin-2 (Prdx2), an antioxidant protein has been recently studied in relation to meat quality parameters such as tenderness (36). These candidate genes discovered in this research which could establish the basis for stable reference biomarkers of lamb phenotype which will be a significant boost for welfare programs.

Collectively, on average ewes that were shorn early in pregnancy demonstrated higher grazing activity, better body condition, and lower stress levels than once shorn ewes. Furthermore, first generation lambs matched to twice shorn ewes expressed visually finer wool with better comfort scores than those F1 lambs that were matched to once shorn ewes. Early shearing of the ewes resulted in improved grazing activity which could have supported the elevated nutritional plane during mid- to late- gestation of the ewes, future research could benefit from the combination of wool cortisol hormone profiling and body condition scores to better evaluate the production characteristics of Merino ewes) (see 37).

Overall, the research outcomes contribute significant new knowledge to the Australian sheep production industry, and it will be a valuable tool for sheep health and welfare assessments in future.

## Materials and Methods

### Ethics approval

This research was originally approved by Western Sydney University ACEC Protocol (A12610) which was later ratified by The University of Queensland ACEC Approval Protocol (SAFS/544/19).

### Experimental Design

The study was conducted on a sheep property in Cattai NSW 2756, Australia. All Merino ewes were shorn prior in October 2018. In January 2019, a total of 100 Merino ewes participated in natural joining. Once joined successfully, 48 mixed age (not maiden) Merino ewes were used for this experiment (the experimental ewes were run together with the rest of the dry ewes). Ewes were bought into the pens by the researcher and visually assessed and conditioned scored by the researcher. Upon confirmation that the ewe was sound (not ill, or lame) she proceeded to inclusion in the experiment, if the ewe was unsound she was not included in the research and removed from the experiment at this point. There were 24 ewes per treatment group (once shorn or twice shorn). All ewes were run as a flock and shearing frequency was the only main factor which was different between the two groups. Ewes (and lambs) were fitted with light weight battery powered collar tags attached with tri-axial accelerometer to discriminate between grazing, standing and walking activity (see 37-39).

### On Farm Assessment

#### Scanning of ewes

Ewes were scanned by experienced operator using a ‘walk through’ system which required minimal set up and consisted of the operator’s crate or crush which ewes enter/exit as they were scanned. During each scan, the ultrasound technician reported an ewe as either pregnant or nonpregnant. Pregnancy is characterized by the presence of a fetus(es) with a heartbeat. Ultrasound technicians can also apply an approximation of length of pregnancy from conception to time of scanning – this is known as early and late from time of conception to scanning.

#### Collection of Wool Fibre

Ewes were bought into the pens and visually assessed and conditioned scored by the researcher. As part of the normal shearing regime on farm a sample of wool was collected from the fleece on the top knot (closest to the skin) as this is an area that can be accessed readily on the animal and recently validated for wool cortisol evaluation by the researchers (17, Sawyer et al., 2021). An ear notch tissue was collected from the ewes and lambs using Allflex TSU sampler (Source: https://www.allflex.global/au/product/tsu-applicator/).

### Laboratory Methods

#### Hormone Analysis

The wool cortisol concentration in each sample was determined by colourmetric analysis using polyclonal anticortisol antiserum (R4866 – supplier UC Davies California, USA) diluted in ELISA coating buffer (Carbonate-Bicarbonate Buffer capsule (Sigma C-3041) and 100 mL Milli-Q water, pH 9.6), working dilution 1:15,000. This was followed by reactivity with Horseradish Peroxidase (HRP) conjugated cortisol label (CJM, UC Davies) diluted 1:80,000, and cortisol standards diluted serially (1.56 – 400 pg/well). Nunc Maxi-Sorp™ plates (96 wells) were coated with 50 μL cortisol antibody solution and incubated for a minimum of 12 hours at 4 °C. Standards, including zeros and nsbs (non-specific binding wells), were prepared serially (2-fold) using 200 μL standard working stock and 200 μL assay buffer (39 mM NaH_2_PO_4_H_2_O, 61 mM NaHPO_4_, 15 mM NaCl).

For all assays, 50 μL of standard and (1:10) diluted 90% ethanol extracted wool samples were added to each well, followed by 50 μL of the cortisol HRP. Each plate was loaded in under 10 minutes. Plates were covered with acetate plate sealer and incubated at room temperature for 2 hours. After incubation, plates were washed 4 times using an automated plate washer (ELx50, BioTek™) with phosphate-buffered saline solution (0.05% Tween 20) and then blotted on paper towel to remove any excess wash solution. Substrate buffer was prepared by combining 1 μL 30% H_2_O_2_, 75 μL 1% tetramethylbenzidine (TMB) and 7.425 μL 0.1 M acetate citrate acid buffer, pH 6.0 per plate. The TMB substrate was added to each well that contained a standard sample at 50 μL to generate colour change. The plates were covered with an acetate plate sealer and left to incubate at room temperature for 15 minutes. The reaction was stopped with 50 μL of Stop solution (0.5 M H_2_SO_4_ and Milli-Q water) added to all wells in the Nunc Maxi-Sorp™ plates. To determine hormone concentration in each sample plates were read at 450nm (reference 630nm) on an ELx800 (BioTeck™) microplate reader. Cortisol concentrations were presented as ng/g.

#### Wool Laserscan Assessment

Snippets of raw wool are cut from samples of raw wool through a mini coring machine. The snippets are removed from the minicore and are washed in a solvent hexane for a period of thirty seconds to wash and blend the fibre snippets together. A shot of compressed air is used to dry the snippets. A random sample is removed *via* tweezers from the washed and dried snippets and placed into a dilute suspension in a mixture of isopropanol (propan-2-ol) and water (8% by volume). The suspension of snippets is transported through a measuring cell which is positioned in a beam of laser light. The reduction in intensity of the laser beam as the individual snippets pass through the beam of light, approximately 500 micrometres in diameter, is sensed by a detector and transformed, using a calibration look-up table, into a diameter in micrometres. A computer is used to collect and summarise the individual measurements to give statistics such as mean and standard deviation of fibre diameter for the specimen. The Laserscan assessment was conducted using an approved Laserscan machine by an independent wool company (Chad Wool Dubbo, NSW).

#### Parentage testing

DNA based parentage determination was done by the Neogen® laboratory, Gatton, Queensland. It provides a very fast and efficient parentage testing service for evaluating the animal’s DNA for accurate parentage determination. Neogen’s lab uses Single Nucleotide Polymorphism (SNP) technology to genetically “fingerprint” each animal at more than 100 chromosomal locations. The animal’s unique genetic fingerprint can be evaluated with the fingerprint of its expected parents by utilizing the knowledge that half of the animal’s genetic information was inherited from the sire (father) and the other half from the dam (mother). Discrepancies in the inheritance pattern would suggest that an animal’s reputed parent would likely be incorrect. This SNP technology enables a higher level of accuracy in parentage testing for cattle, dog, sheep, and pigs (Source: https://genomics.neogen.com/en/parentage-testing-products).

#### Molecular Epigenetic Analysis and bioinformatics

Sheep DNA analysis and quality control were performed using Illumina NovaSeq RRBS (Reduced Representative Bisulfite Sequencing) data of a barcoded 100 bp single end run. RRBS library was produced following the Nuggen’s Ovation® RRBS Methyl-Seq System in the Australian Genome Research Facility (AGRF). The primary bioinformatics analysis involved quality control, trimming adapter, contamination and low-quality fragments, customized Nuggen’s adapter trimming, read mapping, customized Nuggen’s post-alignment processing, DMR (differential methylation region) analysis and annotating differentially methylated genes. Briefly, reads were assessed with FastQC v.0.11.8 and trimmed with Trim Galore v0.5.0. Additional trimming was performed using Nugen’s diversity trimming script with default values. Mapping was carried out with Bismark v0.21.0 to methylation-converted Oar3.1 genome. Alignments were performed with Bowtie2 v2.3.4 aligner with default parameters allowing 0 mismatch in a 20 bp seed.

#### DNA Methylation

EdgeR is a package used to detect and quantify differential methylation of digital RRBS data, that is, counts of reads support methylated and non-methylated cytosines for each locus of a given organism. The library sizes were corrected by the average of the total read counts for the methylated and un-methylated libraries in edgeR. A generalised linear model was then used to quantify the differential expression between the groups. P values were adjusted for multiple hypothesis testing using Benjamini-Hochberg. Nearest transcriptional start site (TSS) was annotated to valid methylation loci using nearest TSS function in edgeR. DMR and all analyzed loci table contained the following fields:Venn diagrams show the numbers of co-upregulated and co-downregulated loci between different comparisons including only loci that have a FDR < 0.05 and logFC > 0.

Raw sequencing reads was firstly processed with TrimGalore to remove adapter/primer, low quality fragments. Trimmed reads then went through a second trimming to remove Nugen’s RRBS specific primer. MultiQC report has been provided to show the statistics of clean reads. Note that the duplicate ratio here looks high because reads digested from the same locus of different molecules have the same sequence in RRBS. These reads were barcoded in library preparation in order to distinguish from true PCR replicates.

Clean reads were then mapped to reference genome (Oar3.1 build) and spike in Lambda DNA. PCR duplicates were removed from the alignments in the following de-duplication step and identical reads from different molecules with unique barcodes were retained. MultiQC report has been provided to summarize the mapping results.

Non-conversion rate: Non-methylated Lambda DNA was used as a spike in to estimate the non-conversion rate of bisulfite-treated genomic DNA. The non-conversion rate was computed by dividing the number of reads support methylated cytosine by the total reads for that cytosine on the non-methylated Lambda DNA, as follows:

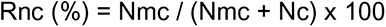

Where Rnc is non-conversion rate, Nmc is the number of reads supporting methylated cytosine and Nc is the number of reads supporting non-methylated cytosine.

#### Differentially methylated regions

In this workflow, one of the most popular Bioconductor packages edgeR pipeline is used for assessing differential methylation regions (DMRs) in RRBS data. It is based on the negative binomial (NB) distribution and it models the variation between biological replicates through the NB dispersion parameter. The analysis was restricted to CpG sites that have enough coverage for the methylation level to be measurable in a meaningful way at that site. We require a CpG site to have a total count (both methylated and unmethylated) of at least 10 in every sample before it is considered in the analysis. The number of CpGs kept in the analysis is ensured large enough for our purposes after filtering. edgeR linear models are used to fit the total read count (methylated plus unmethylated) at each genomic locus. Differential methylation is assessed by likelihood ratio tests, so here we consider FDR value of less than 0.05 as significant DMR.

#### Statistical analysis of other field and lab parameters (Hormone, Body Condition and Grazing Activity)

Statistical analysis was done to test the hypothesis that (1) parameters (hormone, body condition or grazing activity) will be significantly varied between the single or twice shorn pregnant merino ewes. Firstly, data checked for homogeneity of variances and were log-transformed prior to analysis. Statistical analysis was done using a One-way ANOVA using sample, date of sampling and sheep number as the factors and log-transformed data as the dependent variables. Level of significance for all statistical analysis was *p* < 0.05.

## Acknowledgments

Funding for this research was obtained from the Australian Wool Innovation (AWI). DF and RS were awarded the Australian Wool Education Trust (AWET) scholarship. EN and GS conceptualized this research and GS was the field research manager. EN conducted the laboratory hormone analysis and supervised the project team. AT provided valuable input during manuscript writing. Molecular analysis and bioinformatics were conducted by the Australian Genome Research Facility.

