## Supplementary figures and images for "Interplay between stress and reproduction: Novel epigenetic markers in response to shearing patterns in Australian Merino sheep (*Ovis aries*)"

### Table 1

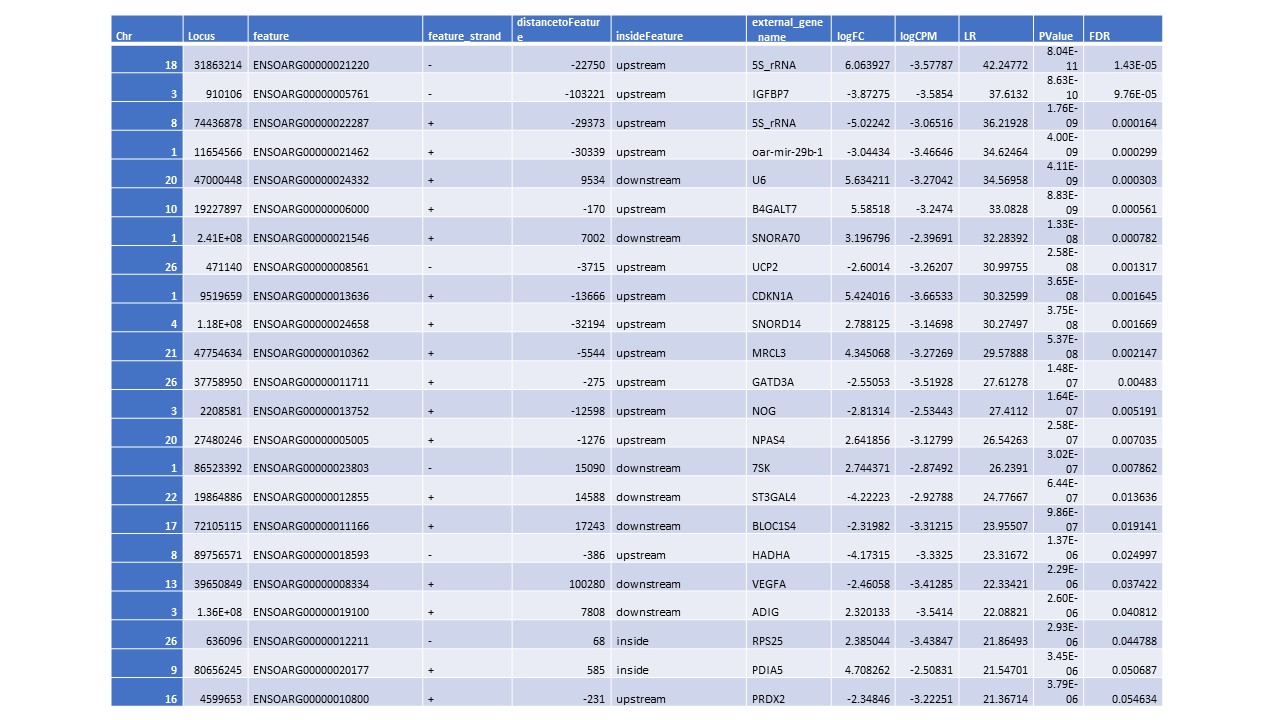

### Table 2

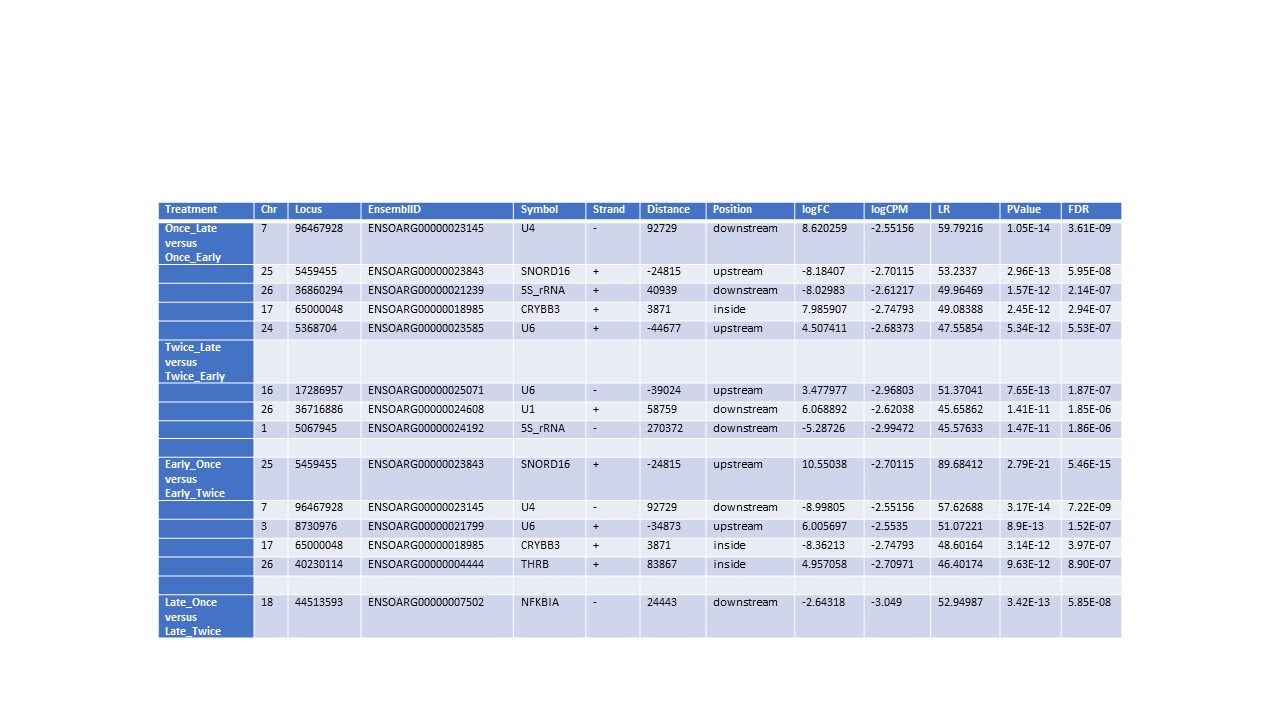

### Table 3

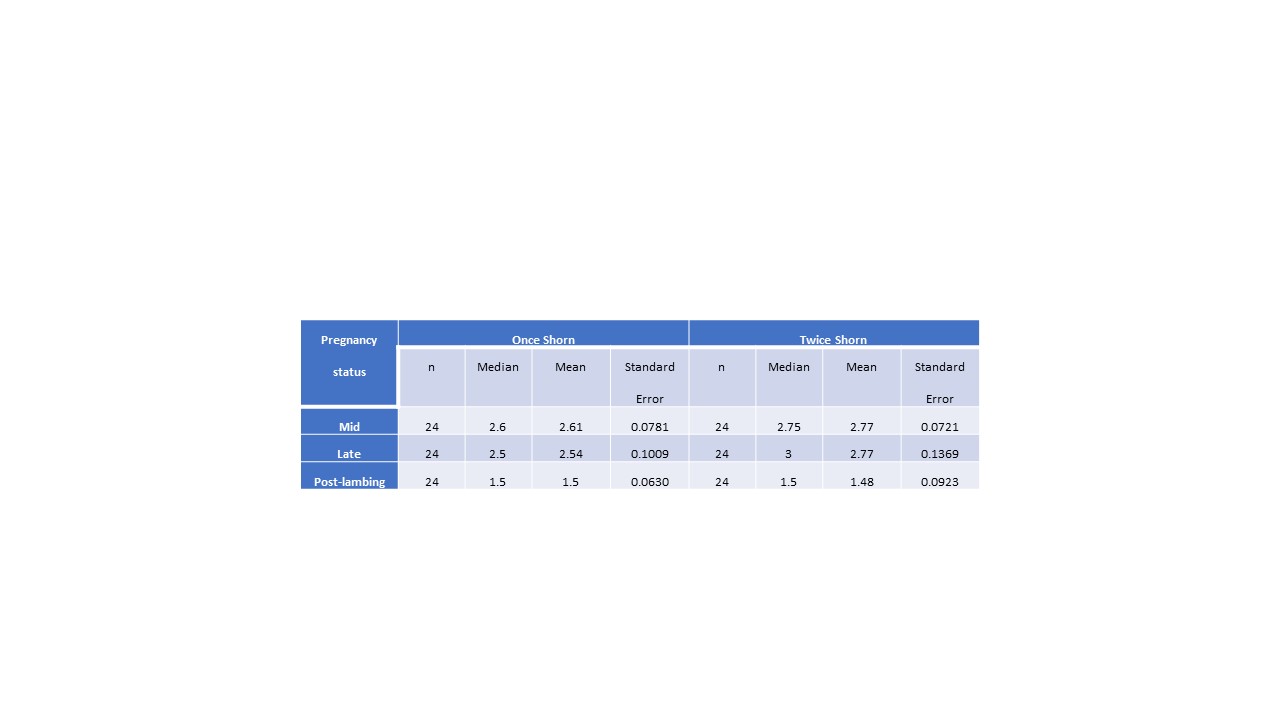

### Table 4

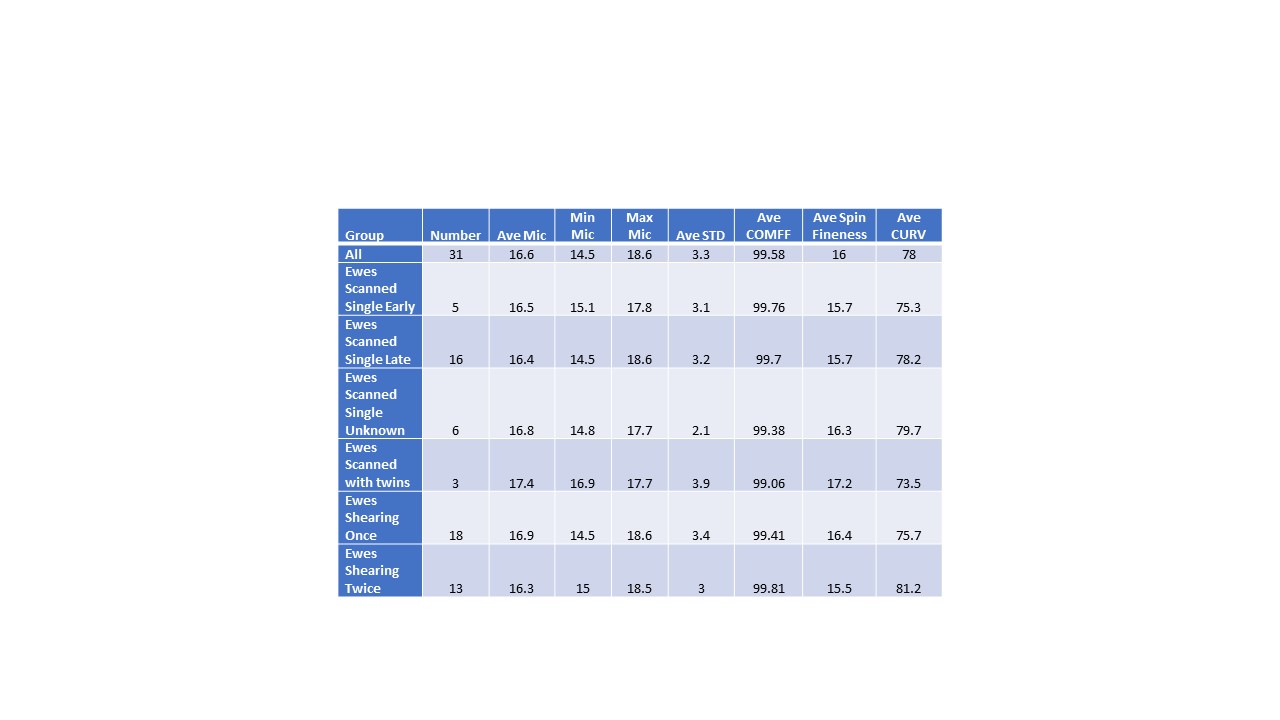
